# Interim impact evaluation of the hepatitis C virus elimination program in Georgia

**DOI:** 10.1101/270579

**Authors:** Josephine Walker, Aaron Lim, Hannah Fraser, Lia Gvinjilia, Liesl Hagan, Tinatin Kuchuloria, Natasha K Martin, Muazzam Nasrullah, Shaun Shadaker, Malvina Aladashvili, Alexander Asatiani, Davit Baliashvili, Maia Butsashvili, Ivdity Chikovani, Irma Khonelidze, Irma Kirtadze, Mark Kuniholm, David Otiashvili, Ketevan Stvilia, Tengiz Tsertsvadze, Matt Hickman, Juliette Morgan, Amiran Gamkrelidze, Valeri Kvaratskhelia, Francisco Averhoff, Peter Vickerman

## Abstract

**Background and Aims:** Georgia has one of the highest hepatitis C virus (HCV) prevalence rates in the world, with >5% of the adult population (~150,000 people) chronically infected. In April 2015, the Georgian government, in collaboration with CDC and other partners, launched a national program to eliminate HCV through scaling up HCV treatment and prevention interventions, with the aim of achieving a 90% reduction in prevalence by 2020. We evaluate the interim impact of the HCV treatment program as of 31 October 2017, and assess the feasibility of achieving the elimination goal by 2020.

**Method:** We developed a dynamic HCV transmission model to capture the current and historical epidemic dynamics of HCV in Georgia, including the main drivers of transmission. Using the 2015 national sero-survey and prior surveys conducted among people who inject drugs (PWID) from 1997-2015, the model was calibrated to data on HCV prevalence by age, gender and PWID status, and the age distribution of PWID. We use the model to project the interim impact of treatment strategies currently being undertaken as part of the ongoing Georgia HCV elimination program, while accounting for treatment failure/loss to follow up, in order to determine whether they are on track to achieving their HCV elimination target by 2020, or whether strategies need to be modified to ensure success.

**Results:** A treatment rate of 2,050 patients/month was required from the beginning of the national program to achieve a 90% reduction in prevalence by the end of 2020, with equal treatment rates of PWID and the general population. From May 2015 to October 2017, 40,420 patients were treated, an average of ~1,350 per month; although the treatment rate has recently declined from a peak of 4,500/month in September 2016 to 2100/month in November-December 2016, and 1000/month in August-October 2017, with a sustained virological response rate (SVR) of 98% per-protocol or 78% intent to treat. The model projects that the treatments undertaken up to October 2017 have reduced adult chronic prevalence by 26% (18-35%) to 3.7% (2.9-5.1%), reduced total incidence by 25% (15-35%), and prevented 1845 (751-3969) new infections and 93 (31-177) HCV-related deaths. If the treatment rate of 1000 patients initiated per month continues, prevalence will have halved by 2020, and reduce by 90% by 2026. In order to reach a 90% reduction by 2020, the treatment rate must increase 3.5-fold to 4000/month.

**Conclusion:** The Georgia HCV elimination program has accomplished an impressive scale up of treatment, which has already impacted on prevalence and incidence, and averted deaths due to HCV. However, extensive scale up is needed to achieve a 90% reduction in prevalence by 2020.

## Introduction

Hepatitis C virus (HCV) causes long-term liver damage and progression to end stage liver disease^1,2^, with deaths due to HCV being greater than malaria in 2015 (~400,000)^3^. An estimated 71 million people are infected world-wide, with 80% concentrated in low and middle income countries (LMIC)^4^. HCV is a highly transmissible blood-borne infection primarily transmitted by injecting drug use and unsafe medical procedures. Until recently, the only treatments available for HCV had poor efficacy, long duration (24-48 weeks) and were poorly tolerated. However, new highly effective all-oral direct-acting antiviral (DAA) treatments are now available, which have made HCV an easily curable infection.

The World Health Organization (WHO) adopted the first global health sector strategy on viral hepatitis (SVH) in 2016, which recognized viral hepatitis as an international public health priority and proposed eliminating it as a major public health threat by 2030^3^. Prior to this, the republic of Georgia, which has one of the highest prevalences of HCV globally (5.4% chronic infection prevalence among adults in 2015^5^), launched the first national HCV elimination program^6^, aiming to reduce HCV prevalence by 90% by 2020.

Georgia has a population of 3.7 million people, with an estimated 150 thousand chronic infections of HCV among adults. Recent advances in treatment for HCV, along with the country’s small population, and political and public support led to the development of a HCV elimination programme for Georgia, supported by Gilead^6^. To help guide the elimination programme, a national serosurvey was conducted in 2015^7^. The serosurvey found heterogeneous levels of HCV infection by gender and age. The highest level of chronic infection (>15%) was amongst men aged 30-49 years, with much lower prevalence rates in females (adult prevalence 2.2%). This heightened HCV transmission amongst men is thought to have occurred during the period of unrest around the collapse of the Soviet Union in 1991, when two civil wars and general economic collapse^8^ resulted in high rates of drug trafficking and injection drug use (IDU) in Georgia^9^. Since then, drug use is thought to have diminished, although recent estimates (2007-2016) still suggest about 2% of adults are people who inject drugs (PWID, 40-52,500)^10–12^, which is high compared to a global average of 0.33%^???^. Transmission is also thought to have been driven by iatrogenic transmission, with the overall quality of medical care and blood transfusion safety remaining low until at least 2009^13^. The age distribution and presumed historical patterns of transmission suggest that the HCV epidemic is in decline, but that a cohort of adults infected 20-30 years ago are likely to be progressing towards advanced liver disease so needing urgent treatment.

We developed a dynamic HCV transmission model to capture the evolving epidemic of HCV in Georgia, incorporating the main drivers of transmission. We use the model to estimate the interim impact of the Georgian HCV elimination program, and then determine whether they are on track to achieving their HCV elimination targets by 2020 or whether strategies need to be modified to ensure success.

## Methods

### Model description and initialisation

We developed a model of HCV transmission incorporating the changing demographics of PWID and the general population in Georgia (Figure 1). The framework of the model used is based on a traditional SI (susceptible-infected) model, because the majority of HCV exposures lead to life-long chronic infection^14^. Curative treatment is incorporated (represented by a compartment T), which if successful leads to individuals becoming susceptible again. The model also includes gender, nine age classes (Figure 1C), and divides all individuals into non-PWID, active PWID, and ex-PWID.

**Figure 1:**
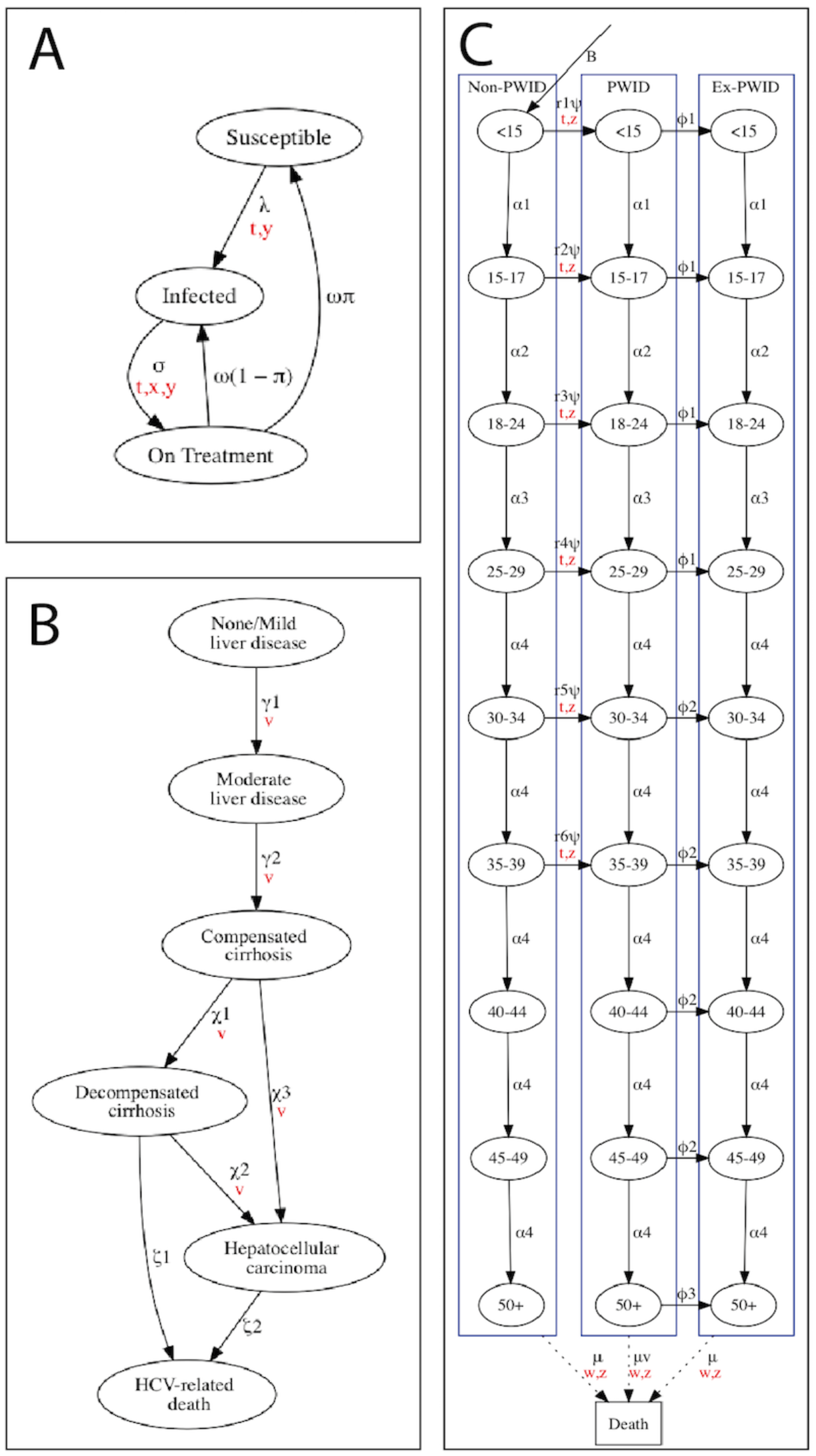
*Flowcharts of state-transitions in the model. A, infection compartments, B, liver disease state compartments, C, PWID and age compartments. Red letters indicate which dimensions the parameters vary with, with t representing time, v representing infection state, w representing age group, x representing liver disease state, y representing PWID state, and z representing sex. Sex compartments are not shown.*

Individuals enter the model in the youngest age group as susceptible non-PWID, equally divided between males and females. Individuals then transition through the age categories, with a proportion transitioning to IDU from all age categories up to age 39, at age and gender-specific recruitment rates. PWID experience drug-related mortality, and cessate from injecting at age-specific rates to become ex-PWID, who die at the same age-specific rates as non-PWID.

Susceptible individuals become infected at a rate that is proportional to HCV prevalence, with a rate of transmission that applies to the whole population, and an additional rate of transmission amongst current PWID. Both these transmission risks are allowed to vary over time to account for changes in intervention coverage in Georgia. The model also allowed for the possibility of assortative ‘like-with-like’ mixing when young (<30 years) and older (>30 years) PWID form transmission contacts, varying between random mixing across these age groups to preferential mixing only between PWID of the same age group.

Upon infection, some individuals spontaneously clear their infection, with the remainder developing chronic infection and gradually progressing through different stages of liver disease (Figure 1B). Individuals with decompensated cirrhosis (DC) or hepatocellular carcinoma (HCC) experience heightened mortality. Treatment occurs at a time-varying rate, with successfully cured individuals returning to the susceptible state with their corresponding level of liver disease, while those failing treatment return to the infected state. After successful treatment, liver disease progression halts for individuals cured with mild or moderate liver disease, while it continues at a slower rate for those with compensated cirrhosis or more progressed disease. Individuals with decompensated cirrhosis or HCC are not eligible for treatment.

The model was initialized in 1900 with a population size of 4 million, all susceptible, non-PWID, and with no liver disease, distributed equally across gender and age compartments. Injecting drug use is assumed to start in 1960. To generate a rapid increase in infection amongst PWID, HCV is seeded in this population with a 10% annual rate of infection for susceptible PWID < 30 years old in the first five years after 1960.

## Model parameterization and calibration

### Calibration and validation data

The model was calibrated to, and compared against available data on the prevalence of HCV from the 2015 National Serosurvey and seven Integrated Biological and Behavioral Surveillance (IBBS) surveys of PWID from 1997-2015 (Table 3). Sero-prevalence estimates from IBBS surveys were converted to chronic prevalence based on the ratio of chronic to antibody prevalence in the National Serosurvey (72%). The National Serosurvey provided gender-specific HCV prevalences in the general population, grouped into three age categories (18-29, 30-49, and 50+), while the IBBS provided year specific HCV prevalence estimates for all PWID, young PWID (18-24) and older PWID (25+). The HCV prevalence estimates used to calibrate the model are given in Table 3, with specific prevalence ratios being used to calibrate the model to ensure it captures increases in HCV prevalence amongst all PWID (16% relative increase over 2006-2015), and the large variations in HCV prevalence by age amongst males. Data from the other IBBS surveys (2001, 2007-2012) were used for model validation. In addition to HCV prevalence data, the model was also calibrated to the overall population size in Georgia in 2015 (estimated from 2014 national census^15^), the estimated number of PWID in 2014 (estimated as a consensus of alternative size estimates^11^), and the proportion of PWID that are 18-29 and 30-49 in the 1998 and 2015 IBBS (Table 3). This last data was included to ensure the model captures the decrease in recruitment of new PWID between these dates.

To compare with our model predictions, we also estimated the observed incidence in PWID between 1997-2001 based on previously unpublished data from a cohort of PWID in Georgia (see supplementary materials).

### Model Parameterisation

The model was parameterised using data from the PWID IBBS surveys, the National Serosurvey, the treatment database for the Georgian elimination program, published literature and WHO databases. All model parameters, uncertainty distributions and their data sources are given in Table 1 and Table 2.

**Table 1:**
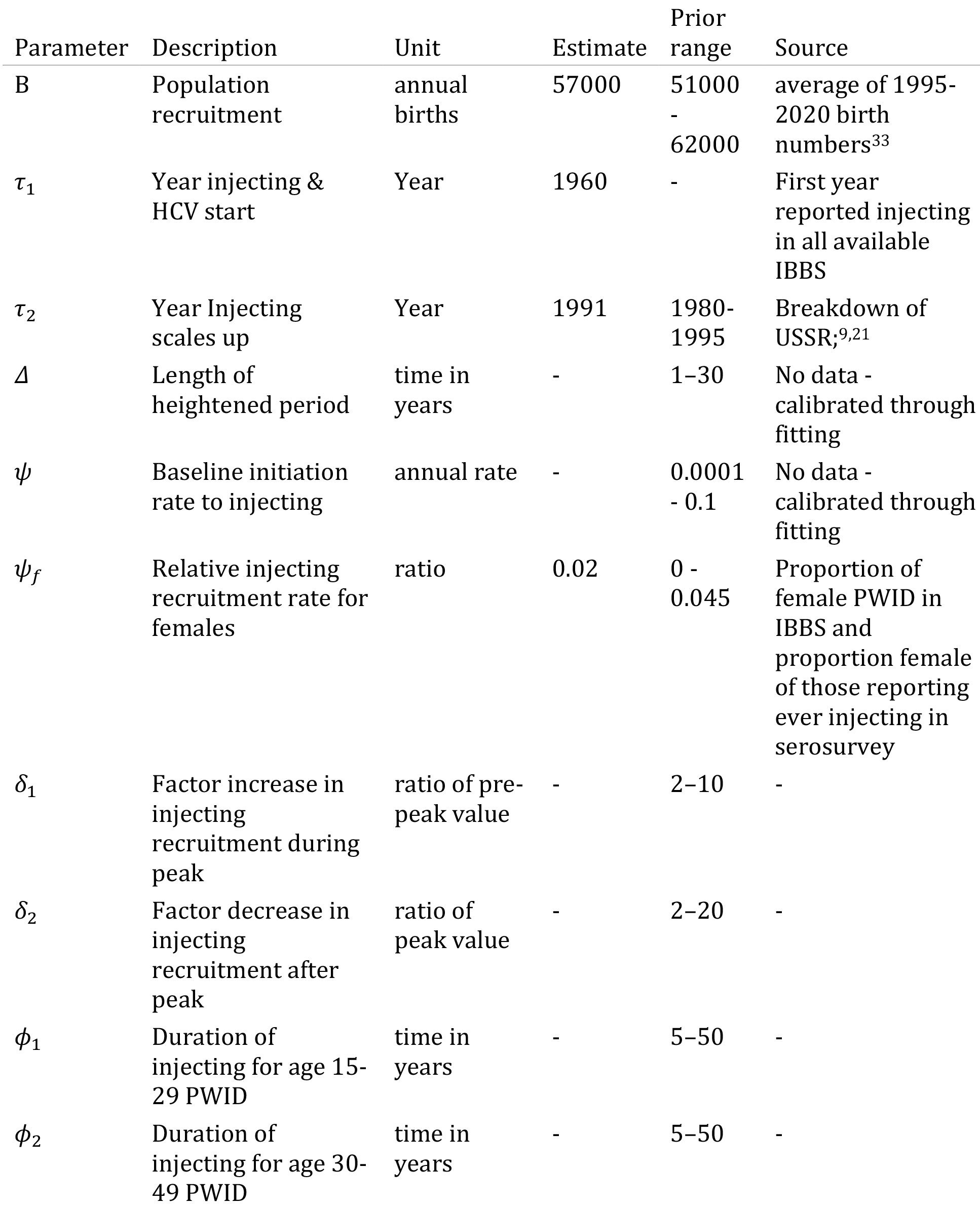
*Parameters varied in model fitting process*

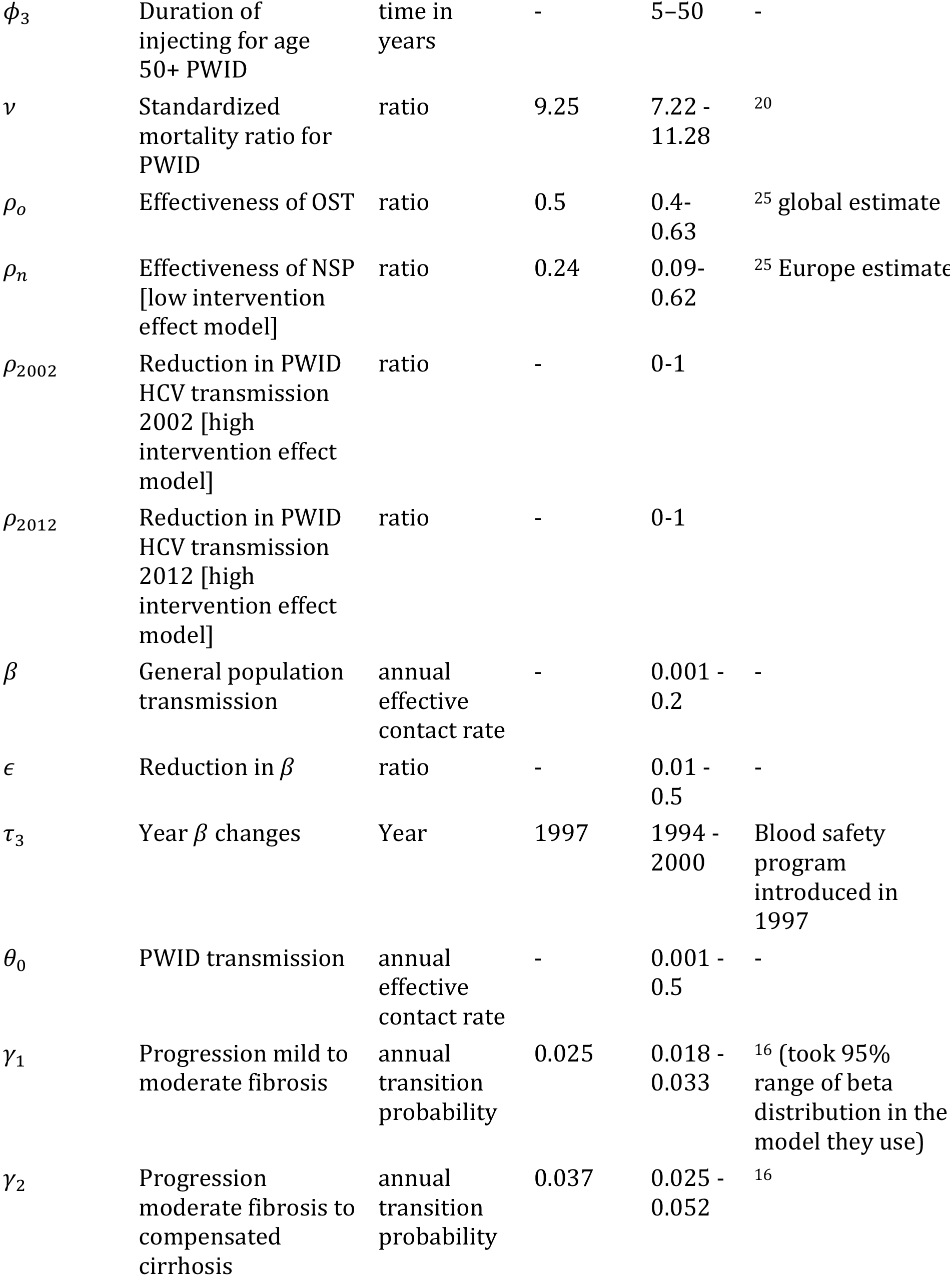

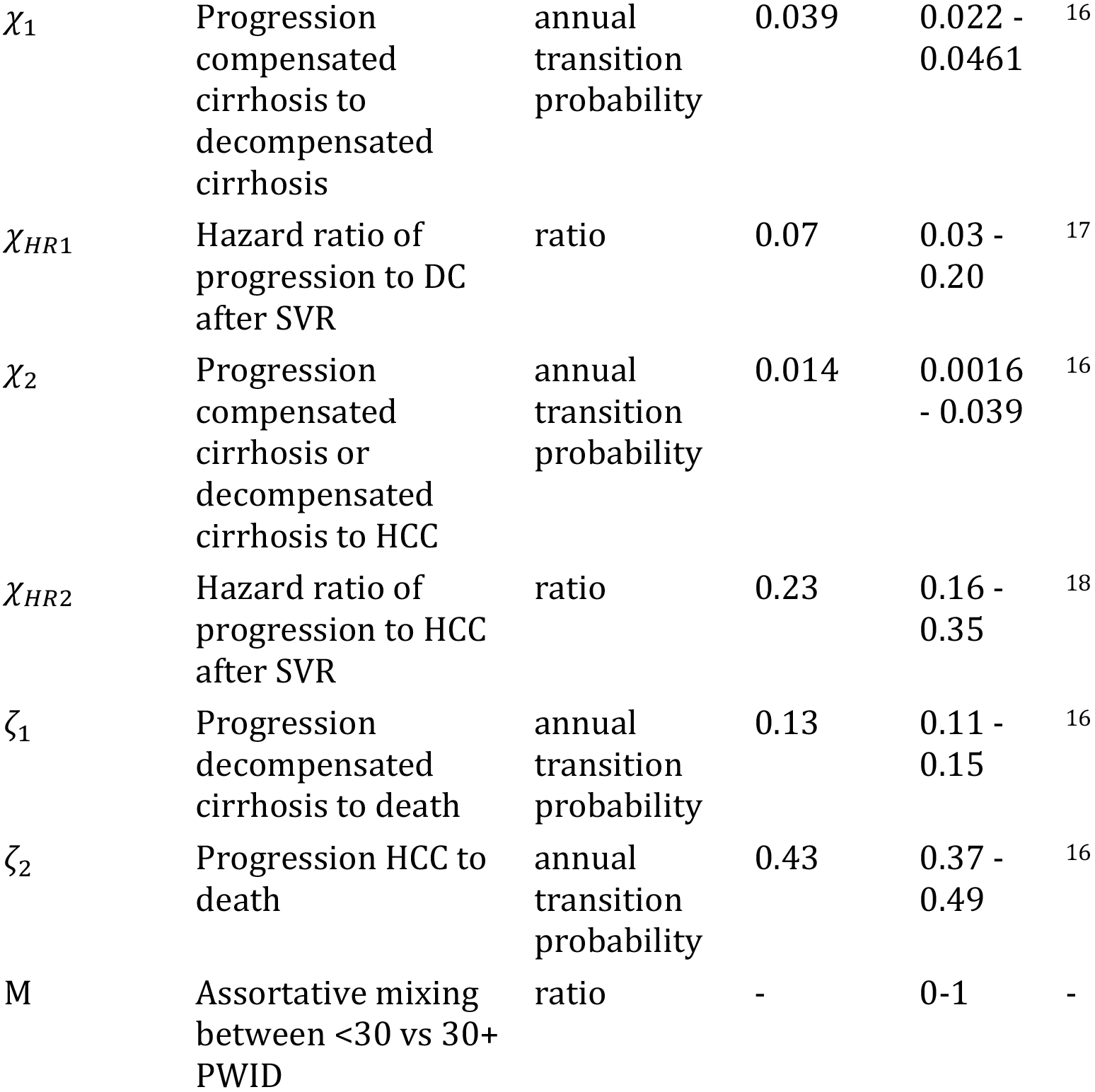

**Table 2:** *Age-varying parameters. Aging rate α inverse of duration of category. Mortality rates from WHO. Relative PWID recruitment from age distribution of PWID (see Supplementary Figure 2).*.

| Age group | $\alpha$ | $\mu_{female}$ | $\mu_{male}$ | PWID recruitment |
| --- | --- | --- | --- | --- |
| <15 | 0.067 | 0.0005 | 0.0005 | $0.02\psi$ |
| 15-17 | 0.333 | 0.0005 | 0.0010 | $0.24\psi$ |
| 18-24 | 0.143 | 0.0005 | 0.0010 | $0.62\psi$ |
| 25-29 | 0.2 | 0.0005 | 0.0010 | $0.09\psi$ |
| 30-34 | 0.2 | 0.0010 | 0.0020 | $0.02\psi$ |
| 35-39 | 0.2 | 0.0010 | 0.0020 | $0.01\psi$ |
| 40-44 | 0.2 | 0.0010 | 0.0040 | 0 |
| 45-49 | 0.2 | 0.0020 | 0.0070 | 0 |
| 50+ | NA | 0.0400 | 0.0700 | 0 |

**Table 3:**
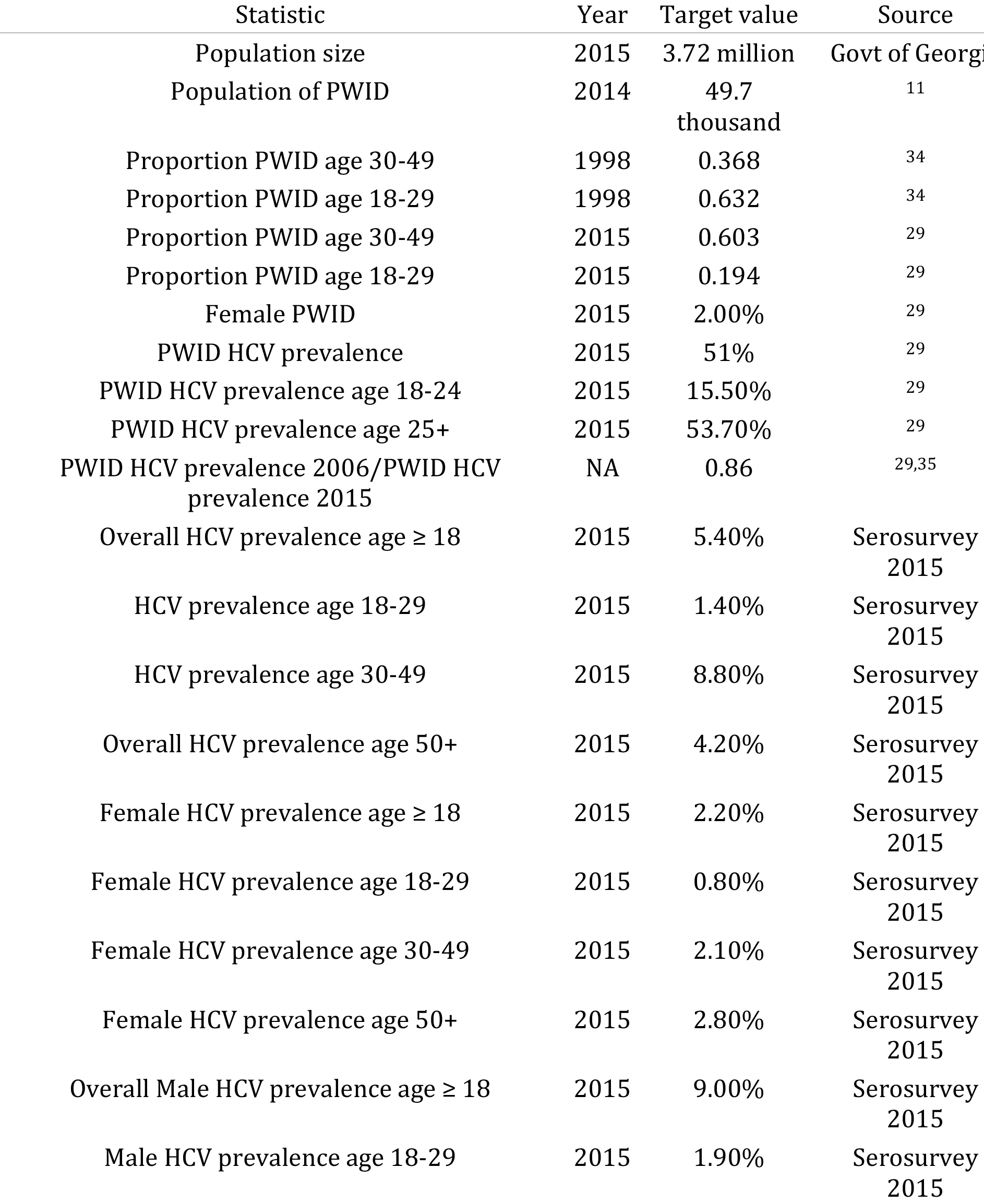
*Summary statistics used to fit model*. \**Ratio of HCV prevalence in young PWID only used in high intervention effect model. Antibody prevalence from PWID serosurveys converted to chronic prevalence at 72% based on chronic prevalence among antibody positive in 2015 general population serosurvey*.

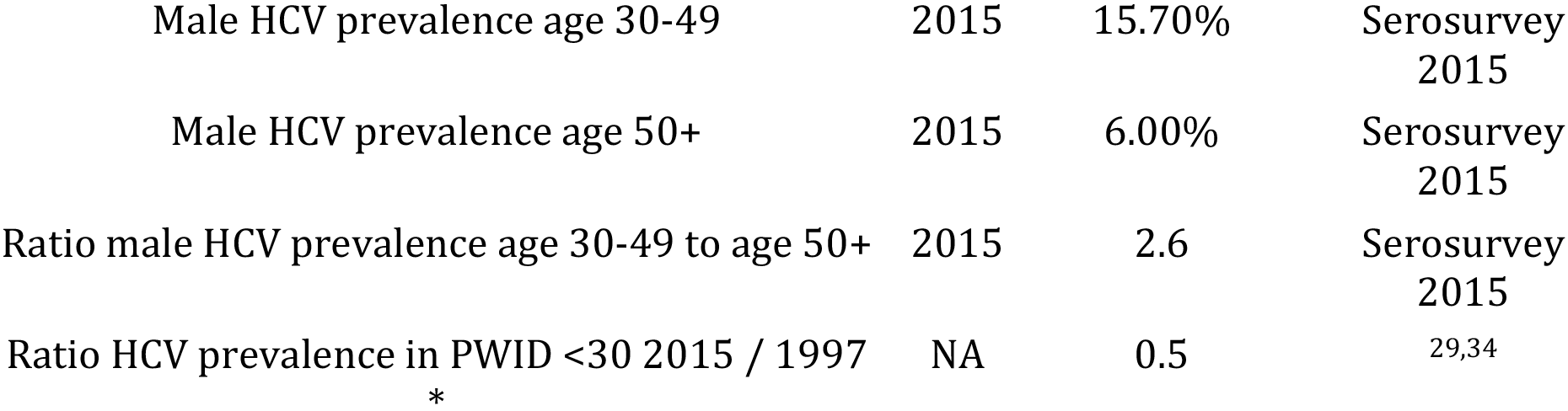

Disease progression and HCV-related death rates were obtained from the literature16–18. Data on HCV treatment rates (see next section) were not used in the model fitting because it began after the last prevalence data used in the model fitting (mid-2015).

Gender- and age-specific mortality rates were obtained from 2015 life tables for Georgia^19^, with PWID having an elevated mortality ratio based on PWID mortality data from Eastern Europe^20^. The model does not account for immigration/emigration or changes in population size.

The number of PWID in Georgia is understood to have increased dramatically alongside the fall of the USSR and resulting social, political, and economic crises^9,21^. Official records of drug users (diagnosed as drug dependent by police) increased eight times between 1990 and 2004, from 2,700 to 21,000, but unfortunately unbiased estimates of the number of PWID over this time period are not available^21^. Recent estimates from 2007-2014 suggest a stable PWID population in Georgia of about 50,000^11^, while IBBS data from 1998-2015 suggest an aging PWID population, likely due to reduced initiation of injecting (Supplementary Figure 3). To account for the likely changing dynamics of IDU in Georgia, we assumed a transient peak in the initiation of IDU, allowing considerable uncertainty in when this occurred and its magnitude (Table 1). Uncertainty also exists around the duration that PWID inject for, which was given wide uncertainty bounds and allowed to vary across age groups. The wide prior ranges for these recruitment and cessation parameters were constrained through fitting the model to IBBS data on the proportion of PWID 18-29 and 30-49 in 1998 and 2015, the estimated population size of PWID in 2014, and the proportion of PWID that are female (from National Serosurvey).

Harm reduction (HR) interventions in the form of needle and syringe provision programs (NSP) was were first introduced in Georgia in 1999, and opioid substitution therapy (OST) was introduced in 2005^22^. Since then, both interventions have scaled up, with 4.5 million syringe kits and 30,330 PWID reached by NSP in 2016 (Georgia Harm Reduction Network, unpublished data) and 4450 PWID on OST in 2015^23,24^. The impact of OST is included in the model by reducing the risk of HCV acquisition (based on a recent Cochrane review^25^) for the proportion of PWID on OST over time^23,24^. However, because uncertainty exists in the impact of NSP in Georgia, we allowed NSP to have greater impact on population-level HCV transmission. This was done to capture the halving in HCV prevalence amongst young PWID (<30 years) in IBBS surveys between 1997 and 2006, which suggests that HCV incidence in PWID, particularly in young/new PWID, may have declined over this period (Supplementary Figure 4). We also undertook a sensitivity analysis where the impact of NSP is used directly from the Cochrane review^25^.

In addition to intervention effects on HCV transmission amongst PWID, the risk of HCV transmission in the general population was allowed to reduced at a point in time to account for other prevention measures, such as the introduction of donor blood screening.

### Model Calibration

We used a modified Markov Chain Monte Carlo Approximate Bayesian Computation (MCMC-ABC) approach to calibrate the model^26,27^ in R version 3.3.2^28^ (see supplementary material). The method obtains a probability distribution of parameter values (the posterior) that constrain the initial prior ranges for model parameters, producing model fits that incorporate the uncertainty in the model parameters and the calibration data. All parameters are simultaneously independently sampled from their prior uncertainty ranges, with the parameter sampling distributions being adjusted iteratively based on how well each sampled run agrees with the calibration data. Importantly, this includes highly uncertain parameters, such as transmission rates, which will be narrowed down based on fitting the model to data.

The parameter sets identified through MCMC-ABC were further filtered to only retain those that agreed (lay in 95% confidence intervals) with the overall HCV prevalence (4.51 − 6.32%) and total female HCV prevalence (1.55 − 2.86%) from the 2015 National Serosurvey, and the HCV prevalence amongst PWID from the 2015 IBBS (45.5 − 56.1%)^29^. These filtered model runs were denoted as the **baseline model fits**.

### Intervention analyses

The baseline model fits were firstly used to estimate the interim impact of the existing scale-up in treatment from May 2015 to October 2017 in Georgia. This utilised data from the treatment program on the number of infected individuals initiating treatment, including treatments targeted to patients with cirrhosis before June 2016 (Table 4). Before March 2016, sofosbuvir with ribavirin was used, achieving per-protocol SVR (sustained viral response) of 80.4%, while after this, ledipasvir with sofosbuvir (Harvoni, Gilead) was used, resulting in an SVR of 98.5% (Georgia Ministry of Labor, Health, and Social Affairs [MoLHSA], unpublished data). We calculated ITT SVR rates for pre-cirrhotic and cirrhotic patients, and an intermediate cure rate taking into account patients who were known to have completed treatment but did not return for SVR testing 12 weeks after finishing treatment (see supplementary materials and Table 4).

**Table 4:**
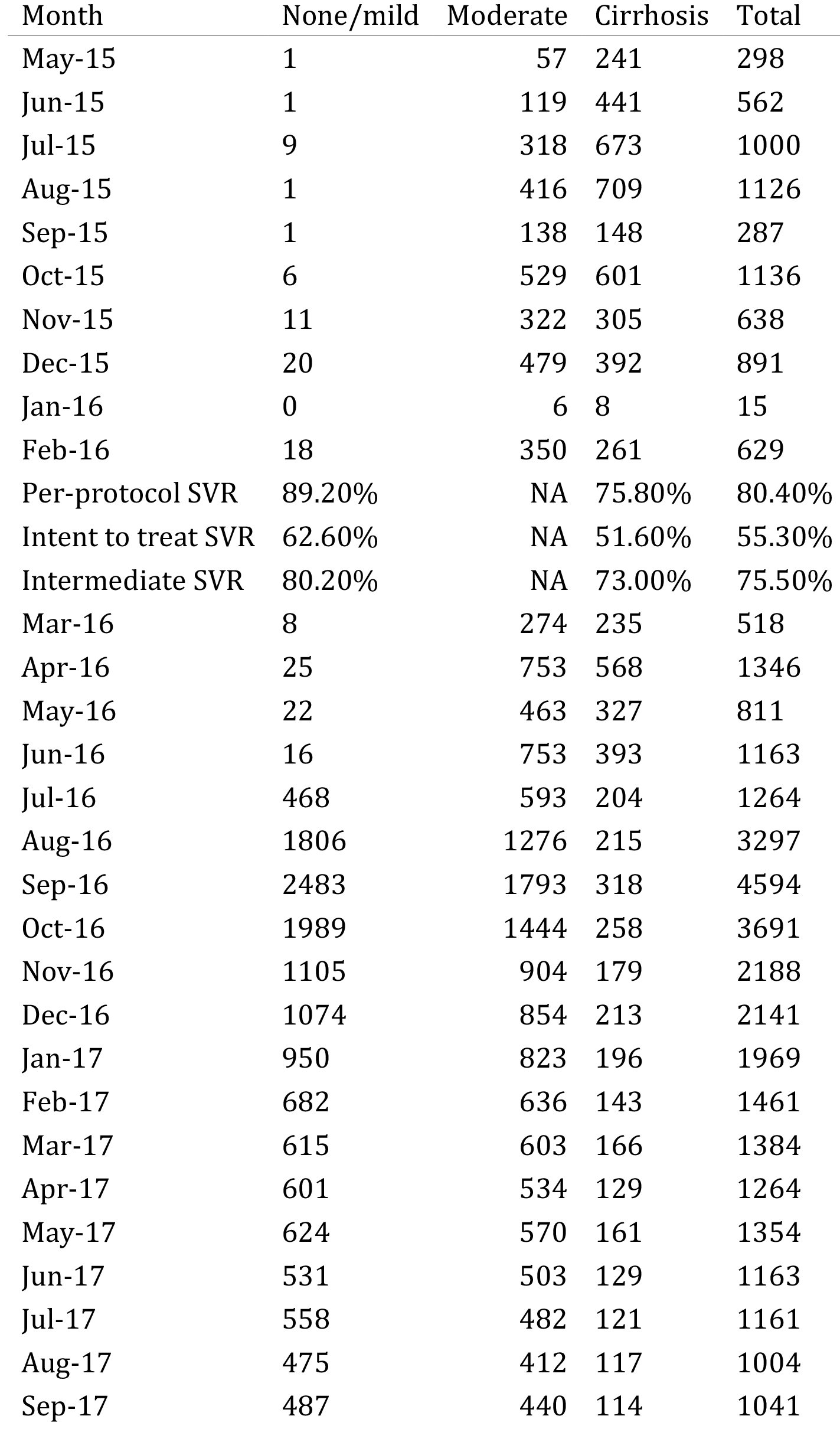
*Treatments by month from data, by liver disease state. Per protocol, intent to treat, and intermediate SVR rates for May 2015-February 2016, and March 2016 to October 2017 are also shown*

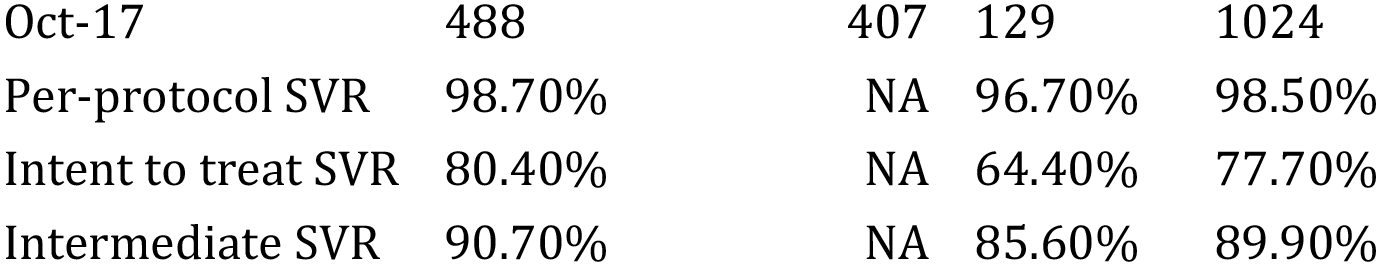

Due to data limitations, there was uncertainty over the number of PWID that were treated. In the base case, we assumed PWID were treated at the same rate as the rest of the population, and compared these results to scenarios where PWID received negligible treatment, or were treated at double the rate of the rest of the population.

Impact was estimated in terms of the relative decrease in incidence and prevalence from May 2015 to October 2017, as well as the number of deaths and infections averted over this period, compared to if no treatment had occurred. The number of infections and deaths averted were also estimated up to the end of 2030, assuming treatment stopped after October 2017.

Following this, we evaluated the impact of alternative intervention strategies going forward (from November 2017), to assess what is required to ensure the elimination program reduces chronic prevalence by 90% by 2020, compared to prevalence levels in May 2015. We considered different treatment targeting scenarios, with treatments either distributed equally across risk groups and disease stages or alternatively targeted to PWID (at twice the rate of other groups) or not (PWID not being treated), or to cirrhotics (80% of infected individuals with cirrhosis (F4) treated annually). For each scenario, we considered the impact up to 2020 of either maintaining the current treatment rate achieved between August-October 2017 (1000 per month), or scaling up to achieve the Georgian government’s estimate of the number of treatments necessary (128,250) to diagnose 90% of cases and treat 95% of these infections (90-95 target). For each treatment targeting scenario, we lastly estimated the treatment rate required from November 2017 to achieve the Georgian elimination target of reducing prevalence by 90% by December 2020 (compared to January 2015 levels). We also estimated what this would achieve in terms of decreasing incidence, and number of infections and deaths averted by 2020.

### Sensitivity analysis

We undertook a sensitivity analysis to determine how the required treatment rate for achieving a 90% decrease in prevalence by 2020 would change if: harm reduction interventions were also scaled up over this period to 75% coverage for OST and NSP; the treatment program achieved the higher per protocol SVR rates amongst all patients; or if existing harm reduction interventions had a lower impact as estimated by a Cochrane review^25^ (see model parameterization section).

## Results

### Baseline epidemic projections without treatment

After model calibration, 554 parameter sets were retained which fit the observed PWID demographics and HCV prevalence for Georgia (Figure 2, Figure 3, Supplementary Figure 2). Prior and posterior parameter distributions are presented in Supplementary Figure 1.

**Figure 2:**
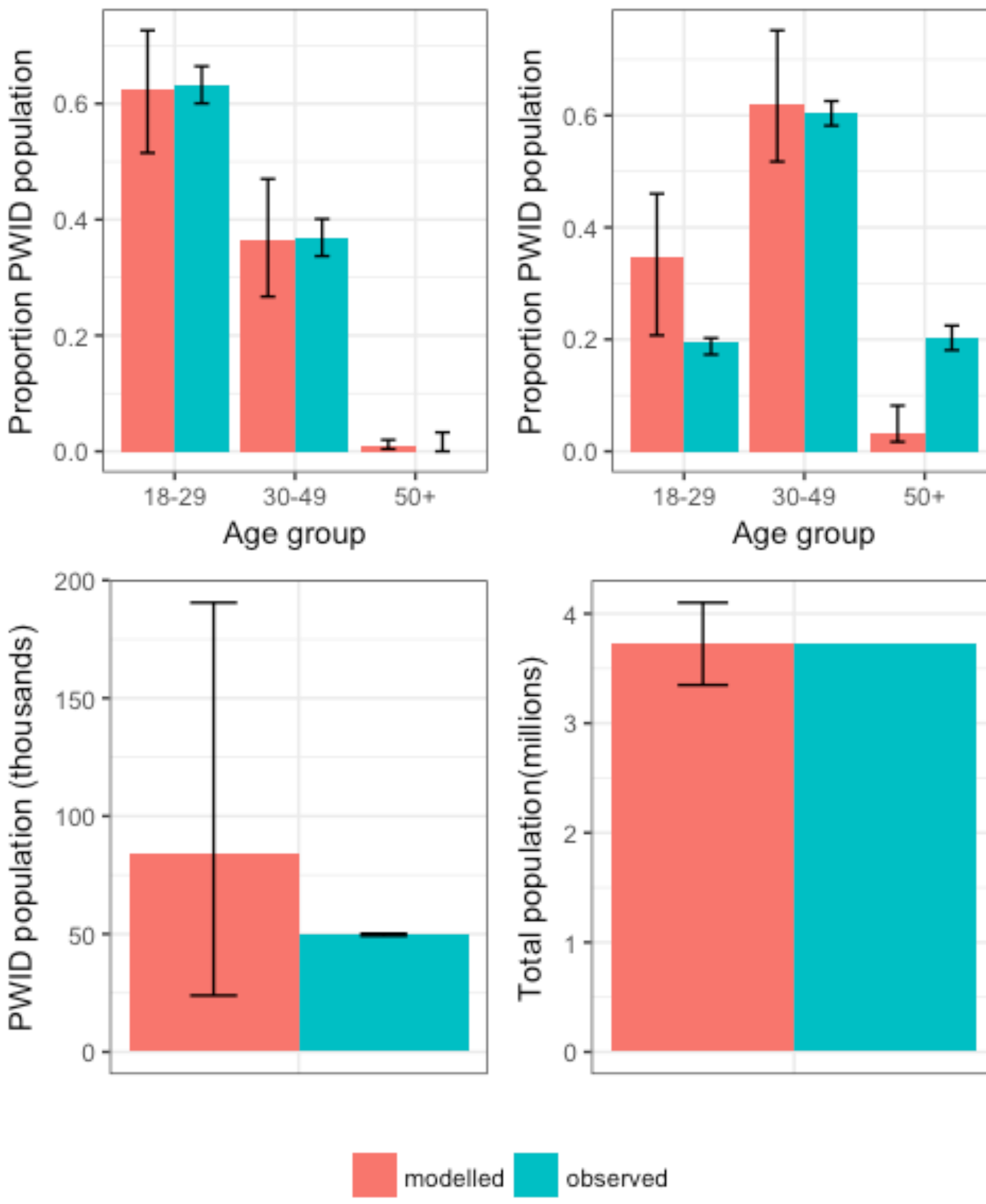
*Model fits to PWID age distributions in 1997 (A) and 2015 (B), and to PWID population size (C) and general population size in Georgia (D)*

**Figure 3:**
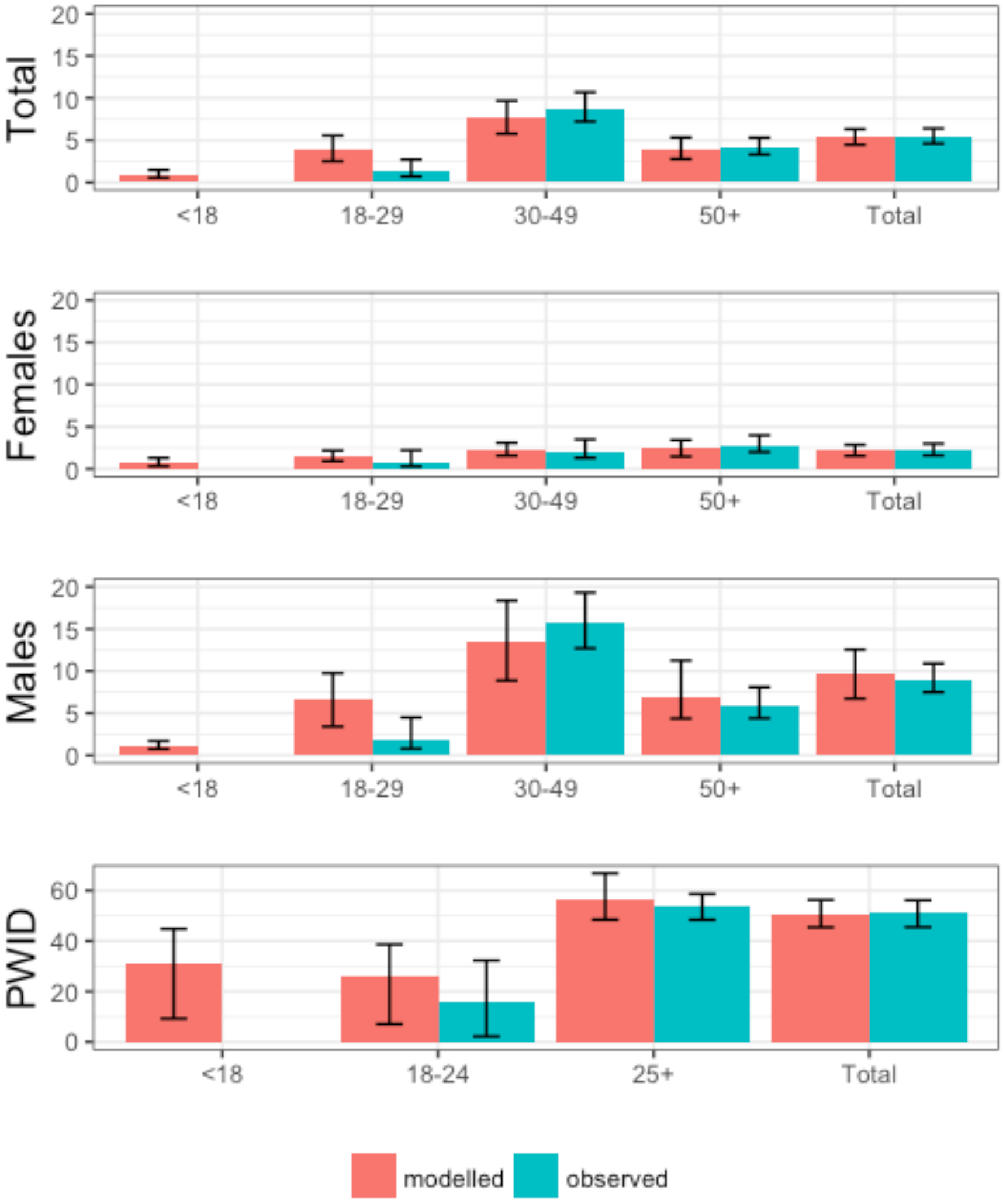
*Fits to percent chronic HCV infection by age and demographic group in 2015, total is total adult population* (≥ 18). *Total, male, and female observed data from national serosurvey, PWID observed data from^29^. No survey data are available for prevalence in individuals <18 years old*.

The baseline model fits suggest that HCV prevalence in PWID is declining over time (Figure 5], middle panel) and HCV incidence in PWID (Figure 6, middle panel) is decreasing or staying stable (percent decrease 10% (−110 − 99% from 2010 to 2015). The population of active PWID is also declining from a peak of 128815 (50583 − 325756) in 2002, with a current population size of 64420 (17598 − 152152). The model is consistent with recent estimates of the adult PWID population, but the modelled estimates for the population of former PWID is more than double the estimate from the 2015 national survey, in which 4.2% (3.5−5.2%) of adults report a history of injecting drug use, indicating a population of 116,760 former PWID (Figure 4). However, this risk factor is likely to be underreported.

**Figure 4:**
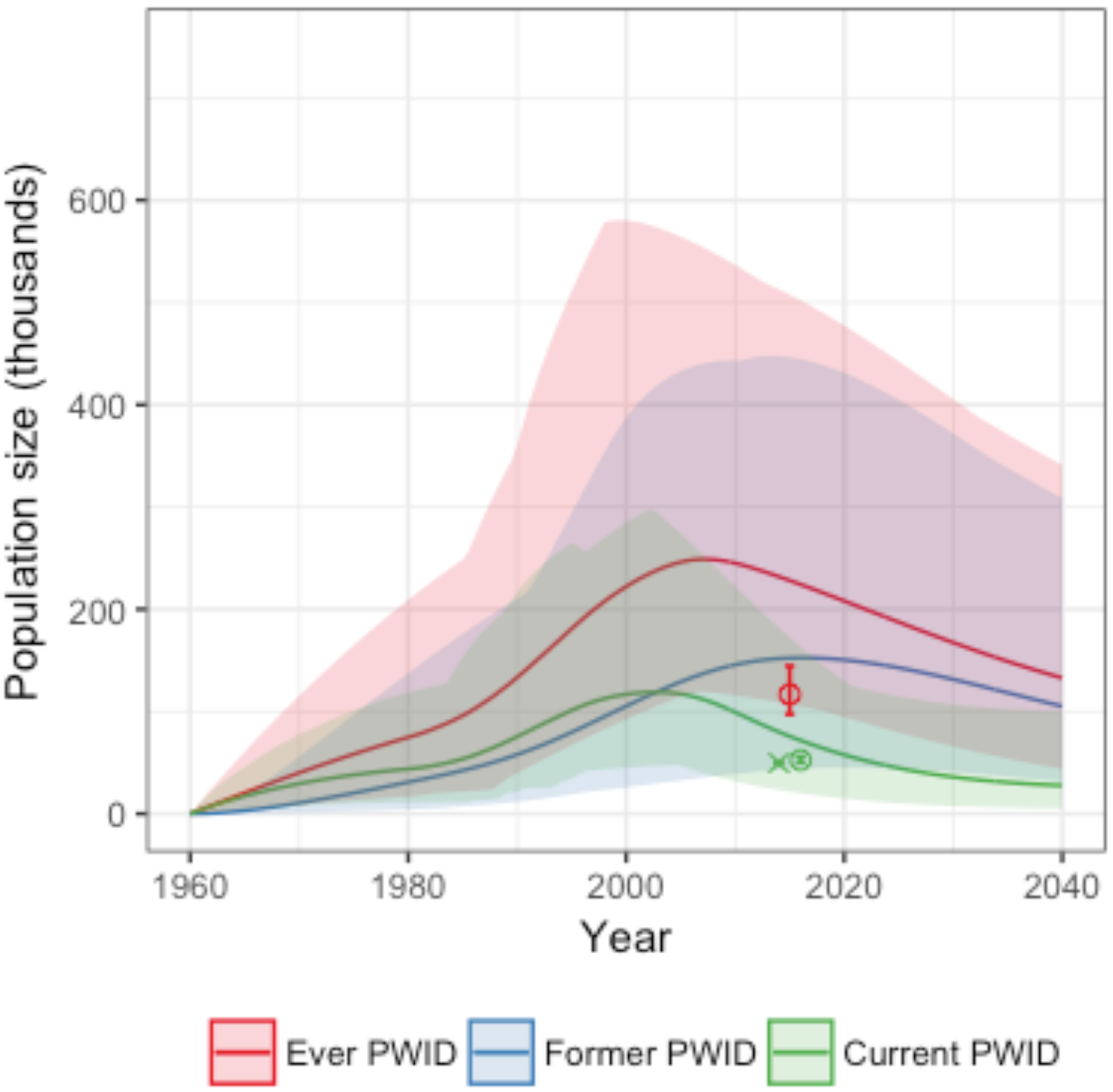
*Projected population size of current, former, and ever (current + former) PWID (adults only) over time. Circles and crosses show available data, with crosses indicating data points used for fitting*.

**Figure 5:**
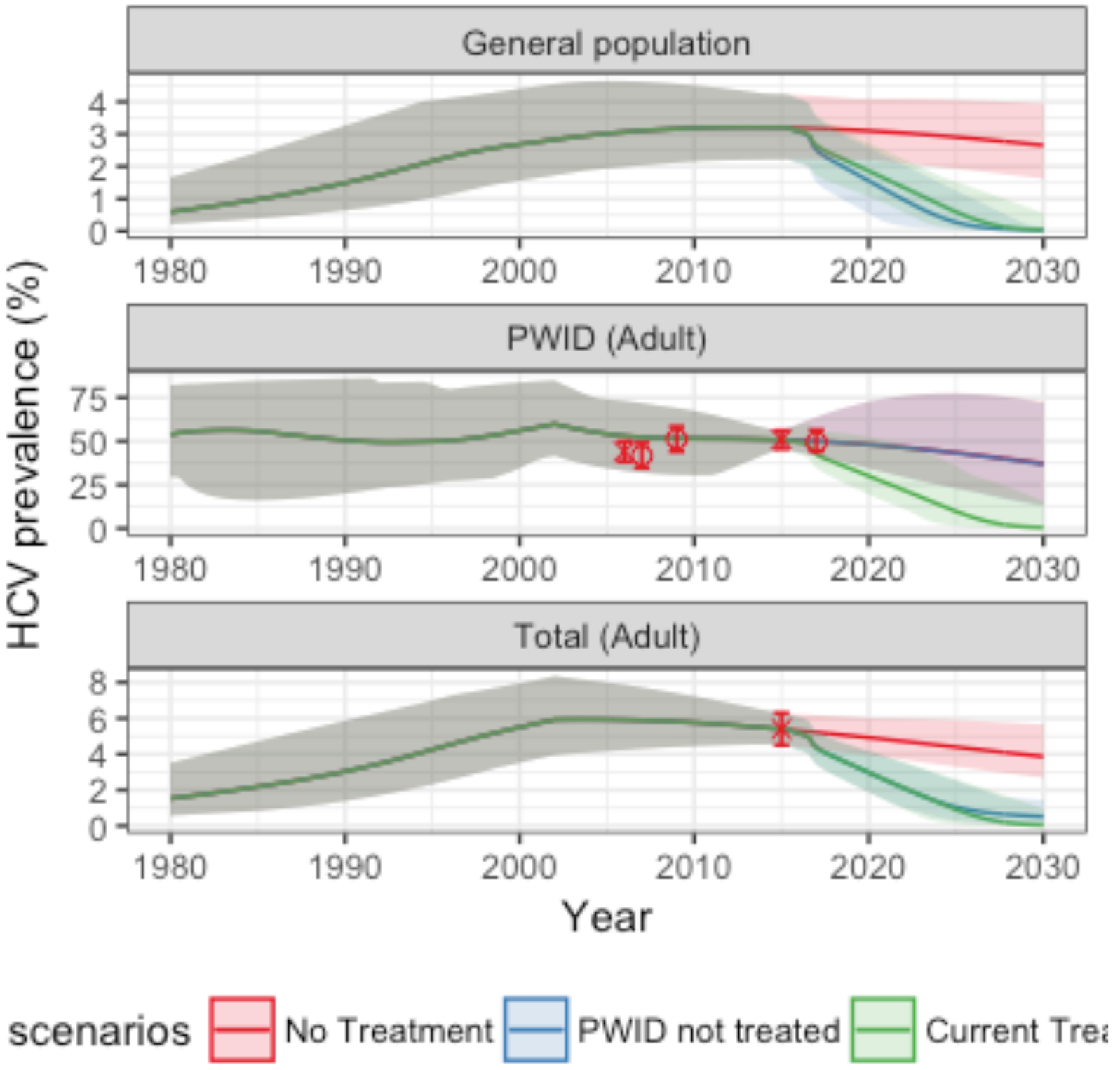
*HCV prevalence over time, in PWID, general population and total. Total and PWID prevalence are for adult only to match available data points, general population is for all ages. Circles and crosses show available data, with crosses indicating data points used for fitting*.

**Figure 6:**
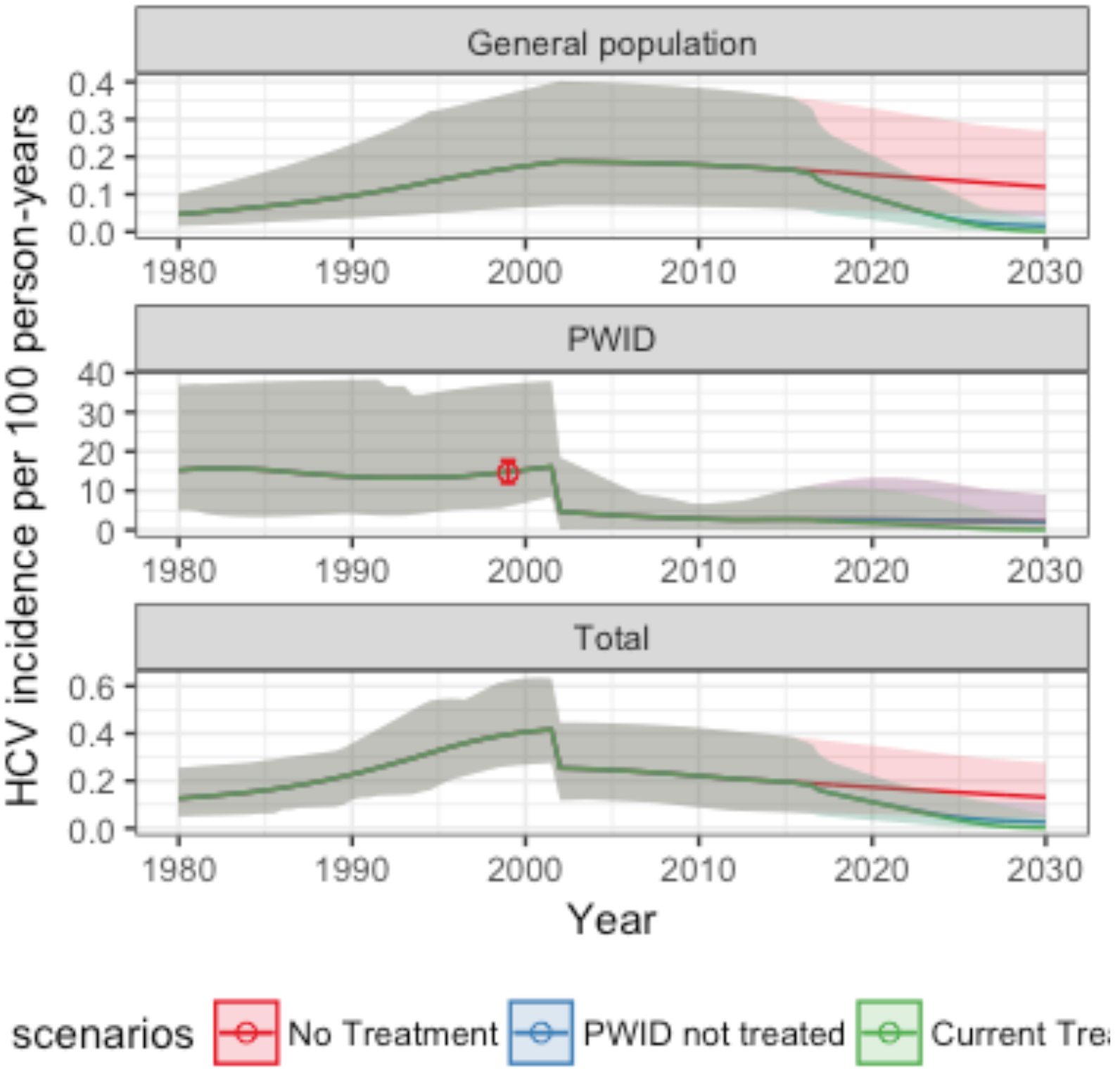
*HCV incidence over time, in PWID, general population and total. Circles show available incidence estimates, these values were not used for model fitting.*

The alternative model calibration estimates a much greater decline in HCV incidence amongst PWID, while the population size of PWID shows less fluctuation than in the main calibration scenario (see supplementary material for results using this model calibration).

With the size of the PWID population and HCV decreasing, the model also projects that the HCV population attributable fraction for injecting drug use (PAF) has decreased dramatically, with PWID being the main drivers of the HCV epidemic in the past, but not now. The HCV proportion attributable fraction (PAF) has declined from 71% (38-92%) over the period 1985-2000 to 37% (13-65%) over 2000-2015 and is projected to be 15% (0-46%) over 2015-2030.

In the absence of any treatment intervention, the model predicts that overall incidence and prevalence will decline from 2015 to 2030 (Figure 5, bottom panel and Figure 6, bottom panel). Incidence will decline in the general population from 0.2 (0.07 − 0.39) infections per 100 person-years in 2015 to 0.13 (0.04 − 0.27) in 2030. In PWID, incidence changes from 2.69 (0.07 − 10.04) to 1.99 (0 − 8.63) over the same time period. HCV related mortality would increase from 607 (165 − 1159) in 2015 to 711 (265 − 1251) in 2030. Annual new infections would decrease from 7009 (2324 − 13214) in 2015 to 4620 (1439 − 10358) in 2030. Total prevalence (all age groups) would decline from 4.2 (3.51 − 4.92) in 2015 to 2.91 (1.94 − 4.43) in 2030.

From the beginning of the program in May 2015, a treatment rate of at least 2050/month would have been required to reach a 90% reduction in prevalence by the end of 2020. In this time period, 40,420 patients were treated, an average of ~1,350 per month. The treatment rate declined from a peak of 4,500/month in September 2016 to 2100/month in November-December 2016, and 1000/month in August-October 2017.

### Interim impact assessment

The 40,420 treatments given are predicted to avert 2654 (1134 − 4419) deaths due to HCV and 16035 (5592 − 37026) new HCV infections by 2030. As of November 2017, the treatment program has averted 98 (31-167) deaths due to HCV and 1517 (596-3585) new HCV infections.

Based on this treatment rate, the HCV adult prevalence is estimated to be 3.91% (2.87 − 4.85%) in November 2017, a decrease of 28% (18 − 37%) since the introduction of the program in 2015. Similarly, HCV incidence has declined by 27% (16 − 37%) from 0.19 (0.07 − 0.39) per 100 person-years in 2015 to 0.14 (0.05 − 0.27) in November 2017.

The above estimates assume that PWID have been reached for treatment at the same rate as the general population. If no PWID have been reached for treatment, the overall HCV adult prevalence and HCV incidence have similarly decreased by 28% (18 − 37%) and 27% (17 − 36%). However, while if PWID are equally treated the prevalence in PWID will reduce by 25% (1 − 39%) in this time period and incidence associated with injecting drug use will reduce by 24% (−10 − 38%), if PWID are not treated, prevalence in PWID will have a percent reduction of 3% (−23 − 17%). Incidence due to injecting drug use will reduce by 4% (−24 − 20%). As of now, the prevalence in PWID would be 49% (39 − 67%) and incidence due to injecting drug use would be 2.28 (0.02 − 12.43).

### Ongoing impact assessment

At the current treatment rate of 1000 patients/month, a 53% (34 − 69%) reduction in adult prevalence and 52% (32 − 69%) reduction in incidence will be reached by 2020, and a 90% reduction reached during the year 2025 (Figure 8, Figure 10).

**Figure 7:**
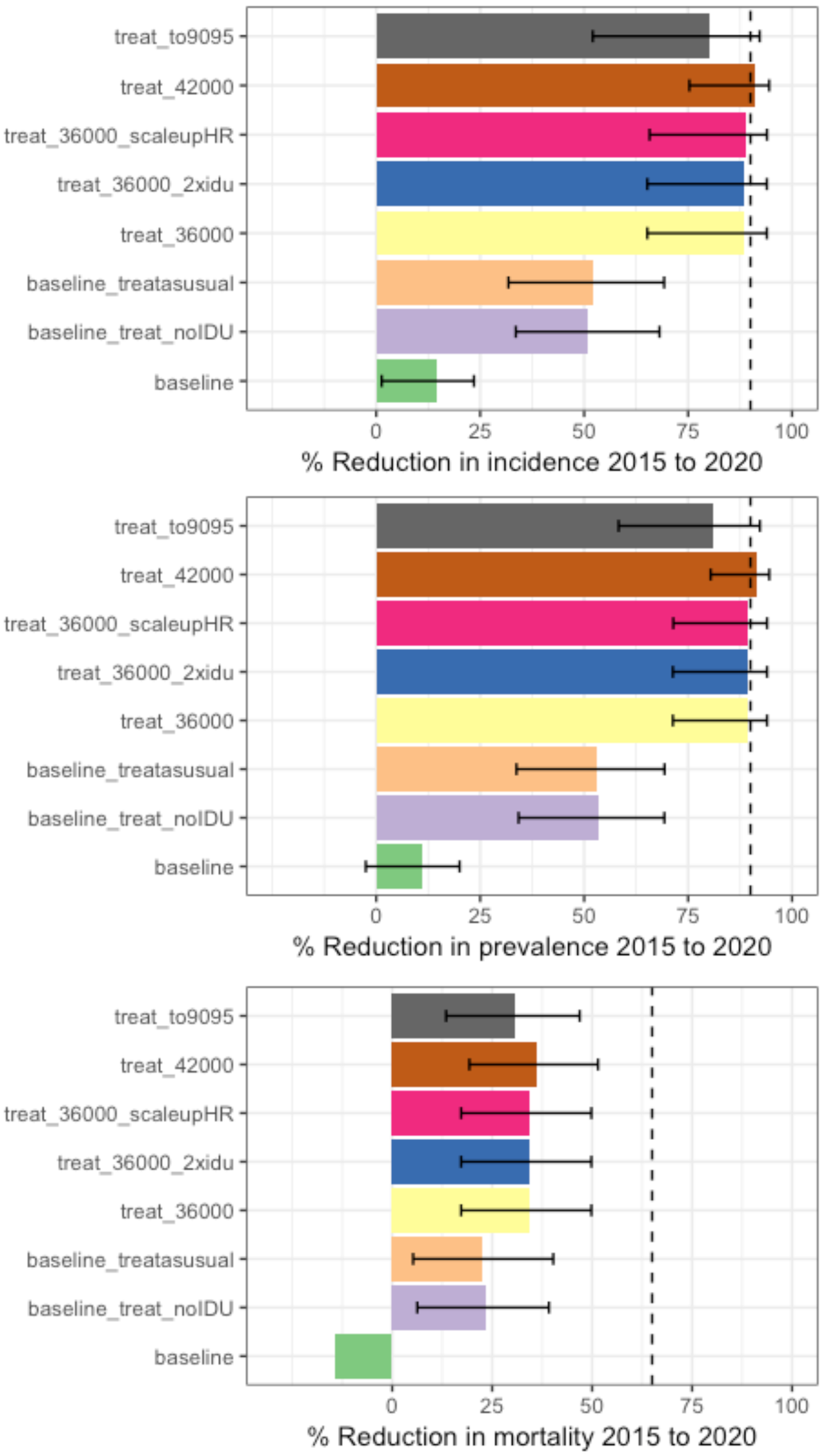
*Percent reduction in incidence, adult prevalence, and mortality over time for selected scenarios. Dashed line shows elimination target of 90% reduction for incidence and prevalence and 65% reduction for mortality*

**Figure 8:**
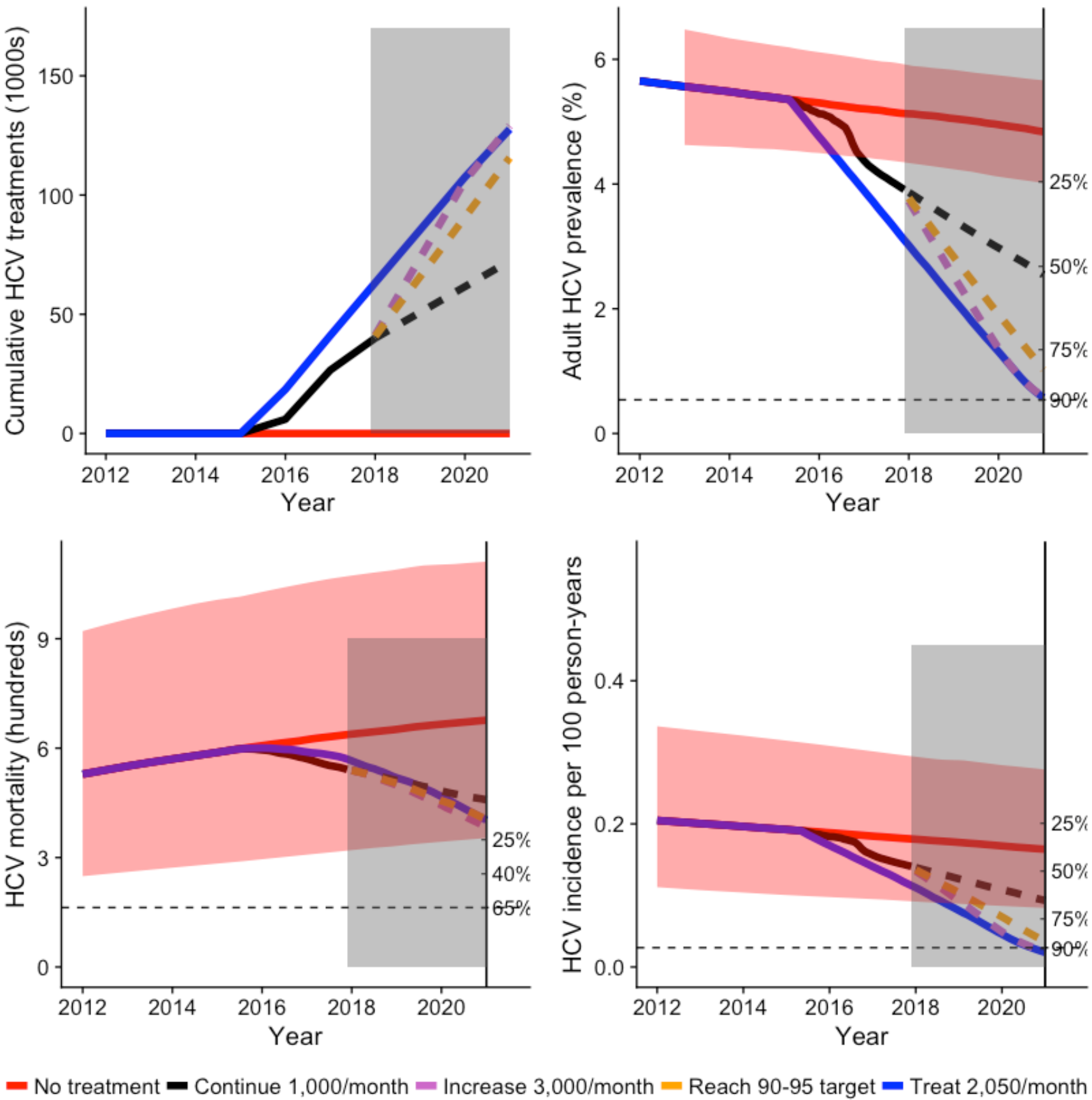
*Interim impact and projected changes in HCV with the treatment program. (A) Cumulative HCV treatments over time, (B) Adult HCV prevalence over time, (C) HCV mortality rate over time, (D) Total HCV incidence over time. The right hand axis shows the % reduction for B,C,D, with horizontal dashed line showing the elimination target. The grey box shows indicates projections into the future. The red line shows no treatment, with uncertainty bounds, the blue line shows the constant treatment rate that would have been required from program initiation to reach the prevalence target by 2020, and the black, purple, and orange lines show existing rates of treatment to October 2017 and projected rates of 1000/month, 2311/month (to reach the 90-95 target), and 3,000/month*

**Figure 10:**
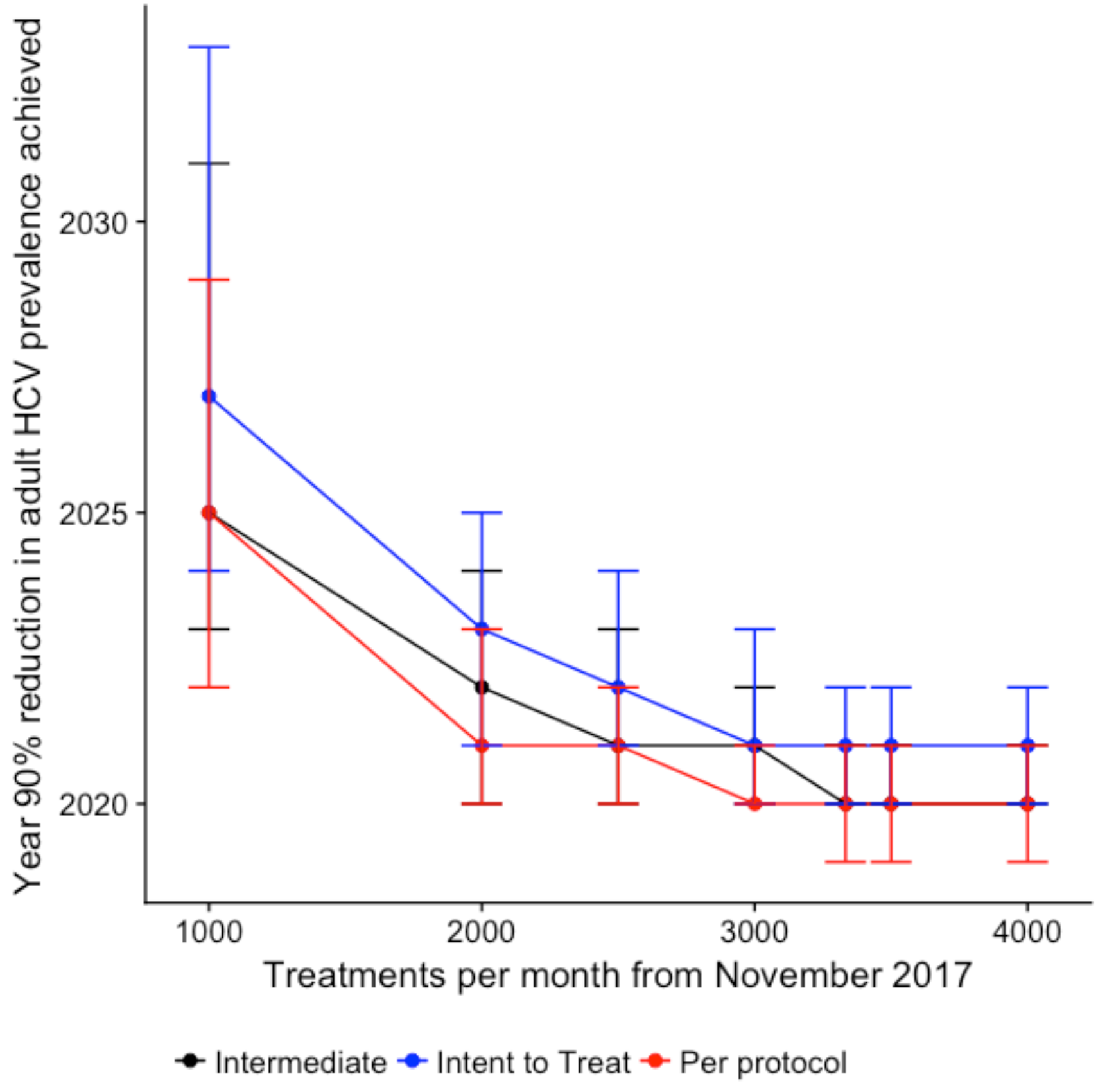
*Year in which 90% prevalence reduction will be reached for levels of treatment scale up from November 2017, with alternative SVR rates: per protocol, intent to treat, and intermediate.*

Scaling up to reach the 90-95 target will achieve a 81% (58 − 92%) reduction in adult prevalence and 80% (52 − 92%) reduction in incidence by 2020, and a 90% reduction reached in 2021.

Reaching a 90% reduction in prevalence and incidence by 2020 will require scale up to 3500 treatments/month with the intermediate SVR estimate. The upper bound (based on intent to treat SVR) requires a treatment rate of 4000/month, while the lower bound (per protocol SVR) achieves a 90% reduction by 2020 with 2500 treatments/month (Figure 10).

Scaling up harm reduction increases the reduction in incidence, while targeting PWID at double the rate of the rest of the population increases the lower bound of the prevalence reduction that will be achieved by 2020 (Figure 7).

Although HCV mortality declines rapidly with the intervention it will not achieve a 65% reduction by 2020. Targeting patients with cirrhosis by treating 80% of cirrhotic patients each you increases the reduction in mortality but it still does not achieve the goal by 2020 (Figure 7, bottom panel). In the first phase of the program, patients with advanced liver disease were targeted, which improved the achieved mortality reduction compared to equal rates of treatment (Figure 8)

## Discussion

Georgia has implemented an ambitious treatment program which aims to reduce the prevalence of HCV by 90% by 2020, and to be the first country to achieve the HCV elimination goals set out in the WHO SVH. By exploring pathways to elimination in Georgia over the next few years, there are many lessons to be learned for HCV elimination globally.

Not all HCV epidemics are created equal. While in many western countries HCV incidence is dominated by transmission through injecting drug use [sources – Australia etc]. In the United States an opioid epidemic is growing and leading to a growing number of HCV cases in young people. In Pakistan, a growing population and increasing HCV prevalence will require an enormous scale up in treatment to reduce the burden of HCV^30^. In contrast, evidence suggests that the overall HCV epidemic as well as the contribution of PWID to HCV in Georgia is declining. The late stage of the epidemic in Georgia means that reducing the prevalence and incidence of HCV is potentially easier than in other settings. However, because many people have been living with HCV for 20 years or more and have already suffered extensive liver damage, no matter how quickly treatment is scaled up, it will be impossible to avert or delay death to HCV for many and a 65% reduction in mortality is unlikely to be achieved by 2020. Treatment rates must be increased, likely tripled or quadrupled, in order to reach a 90% reduction in prevalence and incidence by 2020. One of the primary limitations of the results presented here is that the model does not account for case-finding, or other barriers to maintaining a constant and high rate of HCV treatment as prevalence declines. In the early stages of the program, many HCV cases had already been identified and patients who were interested in treatment came forward. Infected patients may become harder to find as patients remaining are those that are the least likely to be linked to care. Going forward, it will be necessary to screen a large number of individuals to identify patients who are infected with HCV but do not know.

Furthermore, the end game of elimination will result in an increased pool of susceptible (cured) individuals who are able to be re-infected^31^ if concurrent efforts are not made alongside treatment to reduce transmission through preventive measures. All of these factors will contribute to a reduction in the ratio between the number of individuals cured and the resources expended.

Case-finding and linkage to care may be particularly difficult in PWID and former PWID, and although the PWID contribution to the HCV epidemic has declined over time, reaching active PWID for HCV treatment is essential to reduce prevalence and incidence to the target by 2020 or shortly afterwards. While the elimination program has included a dramatic increase in PWID access to HCV testing and treatment over the past few years, and a program in Tbilisi has demonstrated the feasibility of reaching high rates of HCV treatment success among PWID^32^, barriers to HCV treatment access for PWID include continued stigmatization and criminalization of drug use. This also led to high uncertainty in the number of PWID who have already been reached by the program. The Georgia elimination program must continue to make every effort to reach PWID in order to achieve HCV elimination.

There are several limitations to this model, including the necessity of fitting to limited and uncertain data. To account for this, we have presented two model structures which although they differ in the way that incidence, particularly in PWID, has changed over time, both result in similar conclusions regarding the impact of treatment. Additional ways of structuring the model could be developed as more information comes to light on historical patterns of risk, and ongoing monitoring during elimination may help to reveal the true trajectory of HCV incidence. Other studies could be done to estimate uncertain parameters such as the degree of assortative mixing in PWID, the effectiveness of harm reduction measures in Georgia in particular, or spatial heterogeneity in transmission, for example.

The Government of Georgia and partners have made an admirable commitment to eliminate HCV from the country, and the program they have established includes ongoing monitoring and evaluation. The data that they collect will help to steer the elimination program and lessons learned throughout will likely be transferrable to other countries scaling up interventions for HCV.

